# Direct estimation of the spontaneous mutation rate by short-term mutation accumulation lines in *Chironomus riparius*

**DOI:** 10.1101/086207

**Authors:** Ann-Marie Oppold, Markus Pfenninger

## Abstract

Mutations are the ultimate basis of evolution, yet their occurrence rate is known only for few species. We directly estimated the spontaneous mutation rate and the mutational spectrum in the non-biting midge *C. riparius* with a new approach. Individuals from ten mutation accumulation lines over five generations were deep genome sequenced to count *de novo* mutations (DNMs) that were not present in a pool of F1 individuals, representing parental genotypes. We identified 51 new single site mutations of which 25 were insertions or deletions and 26 single point mutations. This shift in the mutational spectrum compared to other organisms was explained by the high A/T content of the species. We estimated a haploid mutation rate of 2.1 x 10^−9^ (95% confidence interval: 1.4 x 10^−9^ – 3.1 x 10^−9^) which is in the range of recent estimates for other insects and supports the drift barrier hypothesis. We show that accurate mutation rate estimation from a high number of observed mutations is feasible with moderate effort even for non-model species.

## Introduction

Being the ultimate source of genetic variation for evolution to act upon, mutation is certainly among the most important genetic processes. The per generation rate at which spontaneous mutations occur in the genome is the central parameter to estimate the effective population size on recent time scales (Charlesworth 2009) or in the course of population history (Schiffels & Durbin 2014), equilibrium of genomic base composition (Hiroshi Akashi & Eyre-Walker 2012) and divergence times (Ho 2014). Yet, the spontaneous mutation rate (µ) is so difficult to measure directly that it has been rarely estimated up to now. Consequently, only very few eukaryotic direct µ estimates are currently available (Denver *et al.* 2009; Keightley *et al.* 2009; Ossowski *et al.* 2010; Keightley *et al.* 2014a; Keightley *et al.* 2014b; Liu *et al.* 2016; Keith *et al.* 2016), scarcely representing biodiversity. More estimates, in particular of non-model species would be highly desirable because they would shed light on the evolution of µ, its associated ecological and evolutionary circumstances (Lynch 2011) as for example the drift-barrier hypothesis (Lynch *et al.* 2016) and Lewontin’s paradox (Ellegren & Galtier 2016).

Currently, two approaches are applied to directly estimate µ: mutation-accumulation (MA) lines experiments and parent-offspring trios (Keightley *et al.* 2009; Keightley *et al.* 2014a). In the MA lines approach, inbred lines are established and bred over many generations (Mukai & Cockerham 1977). Due to the almost absent effect of selection, all except of the most deleterious mutations become eventually fixed and are thus readily identified and confirmed (Denver *et al.* 2009). However, establishing inbred lines is not possible for all organisms, often require complex logistics to transfer generations, intensive care over long time spans and recessively deleterious mutations will be lost with their respective MA-line. In addition, mutator alleles may become fixed, altering the estimated mutation rate (Haag-Liautard *et al.* 2007). In the trio approach, parents together with their offspring are full genome sequenced (Roach *et al.* 2010). This has the advantage that the observed mutational spectrum includes also recessively deleterious mutations as they appear heterozygously in the offspring. Limitations arise from the large number of offspring needed to be screened for an appreciable number of mutations and the requirement to know the parents (Keightley *et al.* 2014a), which is difficult in some species.

In our estimation of µ in the non-biting midge *Chironomus riparius*, we drew on the advantages of both approaches. We established ten MA-lines over five generations and deep sequenced the genomes of a single individual per MA-line. Because individual parenthood is difficult to determine in the swarm breeding *C. riparius,* we compared these individuals with the pooled full-sibling offspring from the single egg-clutch the MA-lines were established from. This yielded an appreciable number of mutations, allowing an accurate estimation of µ.

## Methods

We used a strain of *C. riparius* that was established from a field population in Western Germany and kept since several decades in various laboratories for genetic and ecotoxicological research (“Laufer population”), from which also the *C. riparius* reference draft genome was sequenced (Oppold *et al.* 2016). Larvae from a single egg-clutch were raised at 20°C under most permissive conditions concerning space and food (OECD 2004) to avoid selection. The offspring (F_1_) was allowed to reproduce. After successful reproduction, the adults of this first generation were collected to produce the reference pool (see below). Twenty of the clutches were used to establish as many mutation accumulation lines. These lines were reared as described above for additional four generations, always bringing only a single egg-clutch into the next generation. In the fifth generation, a single individual from each of ten randomly chosen MA lines was retained for genome sequencing (Figure S1). Siblings of each sequenced individual were kept for experimental mutation confirmation.

Due to swarm fertilisation of females, it is not possible to unequivocally determine the parents of a particular egg-clutch. To obtain a baseline against which to identify DNMs, we pooled 190 individual head capsules of their F_1_ offspring and sequenced their pooled DNA to an expected mean coverage of 60X as 150bp paired-end library on a Illumina HiSeq2500 platform, because the allelic composition of the F_0_ parents should be mirrored in the allele frequency of their offspring.

One female individual of each of the ten MA-line was whole-genome sequenced to an expected mean coverage of 25X on an Illumina HiSeq4000 platform. Library preparation of single individuals was performed with the KAPA HyperPrep Kit (KR0961, KAPA Biosystems) to yield enough DNA. The 150bp paired-end reads were individually adapter clipped and quality trimmed, using Trimmomatic(Bolger *et al.* 2014). The cleaned reads of MA line individuals and the reference pool were then processed with the GATK-pipeline (McKenna *et al.* 2010), i.e. mapped with bwa *mem* (v0.7.10-r789, Li & Durbin 2009) against the reference genome draft (NCBI accession number to be provided), duplicates marked with Picard (v1.119 available at http://picard.sourceforge.net), realignment around indels and recalibration of bases with GATK. The raw reads and bam-files for all individuals and the F1 pool can be found at ENA (Project number PRJEB18039).

We established a multistep pipeline in line with Keightley *et al.* (2014a) to identify potential DNMs and minimise false positives. Initially, we applied variant calling with the GATK UnifiedGenotyper (DePristo *et al.* 2011) individually for each MA-line. The reference pool, reflecting the joint genotype of two diploid parent individuals, was treated alike with the exception that variant calling parameters were set as for a tetraploid individual (see above). We then intersected all ten resulting vcf-files amongst themselves and with the reference pool simultaneously, retaining the unique variants for each MA-line, using bcftool *isec* (v1.3, htslib 1.3, available at https://github.com/samtools/BCFtools).

Raw mutation candidates were then quality filtered following the GATK best practices (McKenna *et al.* 2010). Only candidate positions with an overall quality score (GQ) concerning base calling quality and position in read above 90 were considered. Variants with indication for substantial strand bias (SOR > 4) were removed. Since we wanted to concentrate on point mutations and single base indels, indel length was restricted to two. To assure sufficient coverage on the one hand and minimise the effects of mismapped duplicated, paralogous regions on the other, coverage depth was restricted to a range between 15 and 44 reads. A minimum allele count of 5 was required for the non-reference allele to be retained.

The resulting list of candidate positions was then used to create pileups between the respective individual and the reference pool using samtools *mpileup* (SAMtools utilities version 1.1, Li *et al.* 2009). A custom Python-script screened the reads in the reference pool mapped to this position for presence of the alternative allele in the MA-line individual and, if successful, removed it from the candidate list. All surviving candidate positions were manually curated by visualising each candidate position along with the reference pool in IGV (v2.3.68, Robinson *et al.* 2011). This was necessary, because candidate positions contained several false positives due to paralog mismapping (Keightley *et al.* 2014a), PCR artefacts escaping duplicate removal and wrongly emitted variant calls.

The described approach has been shown to yield negligible false negatives (Keightley *et al.* 2014a). Thirty-seven candidate mutations (73%) were checked for presence in the respective MA-line by designing primers for the mutated region and Sanger-sequencing the resulting PCR products for ten full-siblings of the deep sequenced individual. Mutations occurred up to F_3_ should show in this sample at least once with a probability of 0.9989. Assuming that probability of occurrence is equal in all generations, we expected to confirm only about 80% of all mutations because those occurring in the germline of the F_4_ generation will occur only in a single individual in F_5_ (Table S2).

Sites where a mutation could in principal be called according to our criteria were calculated separately for each individual. Assuming a Poisson distribution, we used a Maximum Likelihood method as described in Haag-Liautard *et al.* (2007) to estimate µ and associated confidence intervals.

## Results

The mean sequence coverage ranged from 23.8x to 30.6x for the ten MA-line individuals and was 69.9x for the reference pool, representing the parents (Table S1). The number of callable sites for the MA line individuals ranged between 113 and 130Mb, covering up to 85% of the high complex regions of the genome. Overall, we identified 51 mutations (range 4-11 per individual) of which 26 were SPMs (range 2-6, Table 1). We attempted to validate 37 of these mutations (19 SNPs and 18 indels) by Sanger sequencing. We could confirm 31 of these (17 SNPs, 14 indels, Table S1). By experimental design (Figure S1), we could principally not confirm mutations arisen in the last generation (Table S2), the proportion of unconfirmed mutations was therefore within expectations as were the proportion of confirmed SNPs and indels. Eight mutations were found to be homozygous at least in some of the individuals that were Sanger-sequenced for confirmation. This is not significantly different from the expected mean as inferred from a simulation (χ² = 0.343, p = 0.56, Table S2). One mutation was in an intron of an annotated gene. This is less than expected from the extent of the gene space (15%) in the draft genome.

**Table 1.**
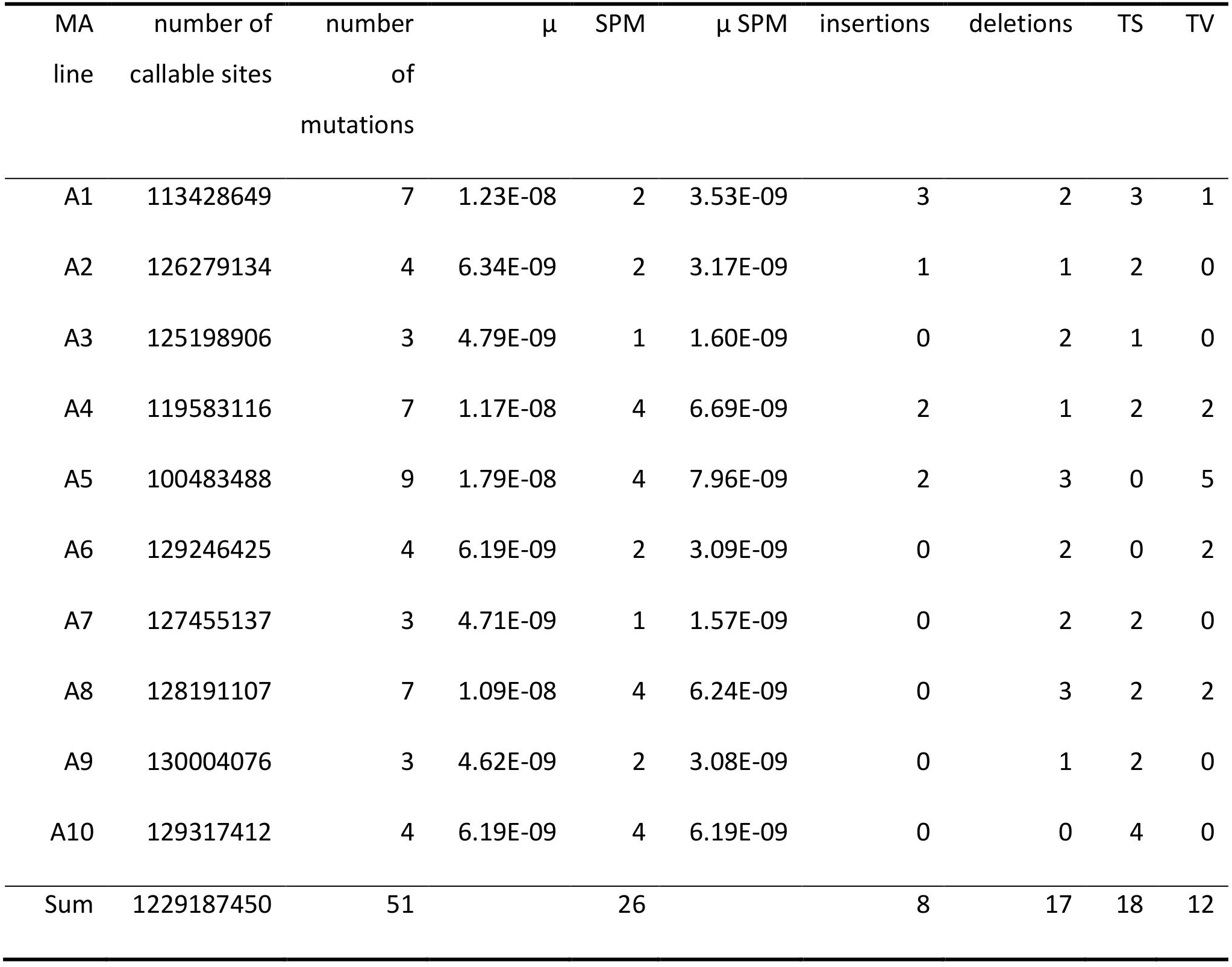
Summary information on the number of callable sites, the total number of single base mutations, resulting mutation rate (µ) per generation and site, the number of single point mutations (SPM) and the associated rate (µ SPM), number of insertions, deletions, transitions (Ts) and transversions (Tv) per mutation accumulation (MA) line.

| MA line | number of callable sites | number of mutations | $\mu$ | SPM | $\mu$ SPM | insertions | deletions | TS | TV |
| --- | --- | --- | --- | --- | --- | --- | --- | --- | --- |
| A1 | 113428649 | 7 | 1.23E-08 | 2 | 3.53E-09 | 3 | 2 | 3 | 1 |
| A2 | 126279134 | 4 | 6.34E-09 | 2 | 3.17E-09 | 1 | 1 | 2 | 0 |
| A3 | 125198906 | 3 | 4.79E-09 | 1 | 1.60E-09 | 0 | 2 | 1 | 0 |
| A4 | 119583116 | 7 | 1.17E-08 | 4 | 6.69E-09 | 2 | 1 | 2 | 2 |
| A5 | 100483488 | 9 | 1.79E-08 | 4 | 7.96E-09 | 2 | 3 | 0 | 5 |
| A6 | 129246425 | 4 | 6.19E-09 | 2 | 3.09E-09 | 0 | 2 | 0 | 2 |
| A7 | 127455137 | 3 | 4.71E-09 | 1 | 1.57E-09 | 0 | 2 | 2 | 0 |
| A8 | 128191107 | 7 | 1.09E-08 | 4 | 6.24E-09 | 0 | 3 | 2 | 2 |
| A9 | 130004076 | 3 | 4.62E-09 | 2 | 3.08E-09 | 0 | 1 | 2 | 0 |
| A10 | 129317412 | 4 | 6.19E-09 | 4 | 6.19E-09 | 0 | 0 | 4 | 0 |
| Sum | 1229187450 | 51 |  | 26 |  | 8 | 17 | 18 | 12 |

Eighteen SPMs were transitions (Ts) and twelve transversions (Tv), resulting in a Ts/Tv ratio of 1.50. There were twice as many G/C to A/T mutations than vice versa (14 to 7), but this difference was not significant (χ² = 2.333, p = 0.13). We observed eight insertions and 17 deletions, a difference that only marginally deviated from random expectations (χ² = 3.240, p = 0.07). With few exceptions, indel mutations were associated with an A or T monomer stretch of at least five positions length (Table S1). Two Sanger confirmed mutations showed different bases in different individuals (A2 scaffold 31: 1191434, G>C and G>A and A7 185:145141, G>T and G>A).

Our estimate of the haploid SPM rate was µ=2.12x10^−9^ (95% CI: 1.39x10^−9^ – 3.03x10^−9^, Figure 1A). Using coalescence estimates of theta from 70 unlinked non-genic regions in European wild populations between 0.023 and 0.034 (unpublished data), we estimate N_e_ for these populations to range between 2.72x10^6^ and 4.02x10^6^. The overall rate for single base deletions was µ_del_ = 1.39x10^−9^ (95% CI = 0.82x10^−9^ – 2.07x10^−9^) and for insertions µ_ins_ = 0.65x10^−9^ (95% CI = 2.85x10^−10^ – 1.22x10^−9^). The total mutation rate for all single base mutations was thus 4.15x10^−9^ (95% CI = 3.13x10^−9^ – 0.54x10^−8^, Figure 1A).

**Figure 1.**
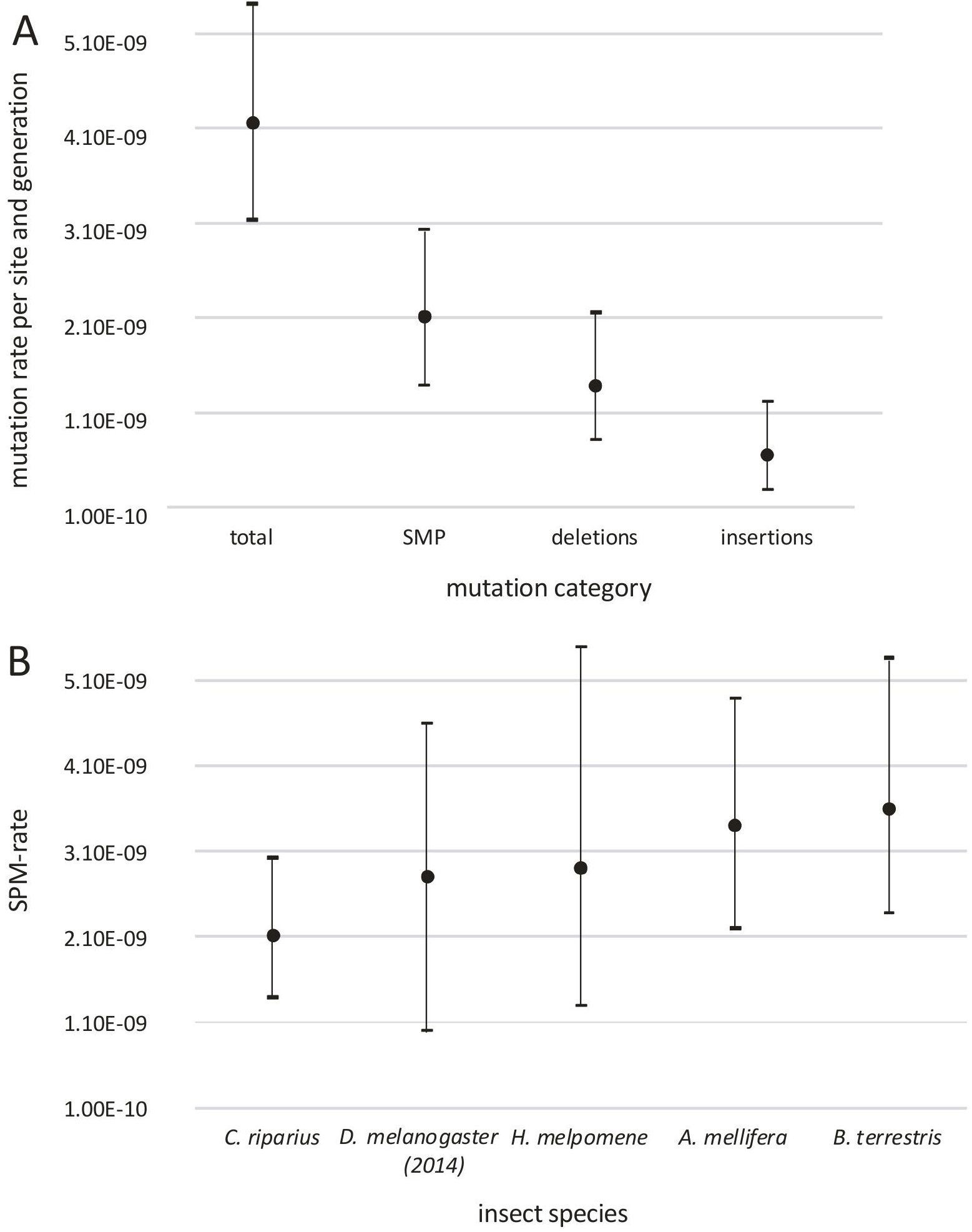
A) Mean and confidence interval estimates for the total single base mutation rate, the single point mutation (SPM) rate, the deletion and insertion rate. B) Comparison of the known SPM rates for insects with confidence intervals where available

## Discussion

We here present a direct estimate of the spontaneous mutation rate in *C. riparius*, a valuable resource from a non-model dipteran as additional representative for the vast biodiversity of insects. We were able to confirm the expected proportion of mutation candidates by Sanger sequencing, suggesting a very low false positive rate. Together with the known low false negative rate of the applied bioinformatics pipeline (Keightley *et al.* 2014a), the presented values likely are accurate estimates. The estimated SPM rate reported here is in the range, although at the lower margin, of estimates from both MA-lines and single generation parent-offspring approaches in *D. melanogaster* (2.8x10^−9^ to 5.49x10^−9^, Keightley *et al.* 2014a; Schrider *et al.* 2013; Keightley *et al.* 2009) or *H. melpomene* (2.9x10^−9^, Keightley *et al.* 2014b), with broadly overlapping confidence intervals. It is however, much lower than the recently reported rates for *A. mellifera* and *B. terrestris* (Yang *et al.* 2015, Liu *et al.* 2016, Figure 1B). The estimate of the *C. riparius* effective population size is comparable to both, *D. melanogaster* and *H. melpomene* (~1.4x10^6^ or ~2x10^6^ for *D. melanogaster* and ~2x10^6^ for *H. melpomene* (Keightley *et al.* 2014a; Keightley *et al.* 2014b)). This and the similarity of µ may be taken as support for the drift-barrier hypothesis (Lynch *et al.* 2016), stating that the realised µ of a species is determined by the balance between selection and drift.

Even though not significantly different, the observed bias towards G/C > A/T mutations is in line with the high A/T content of the *C. riparius* genome (G/C content 31%, Oppold *et al.* 2016). This A/T bias of the *C. riparius* genome is probably also responsible for the observed shift from SNPs to indel mutations compared to other organisms. It leads statistically to an over proportional increase in AT monomer runs and thus to potential mutation sites. Indeed, indel mutations were significantly more often observed in monomer stretches than expected, and their mutation rate increased exponentially with stretch length (Supplement Text 1). Comparison to *D. melanogaster* (G/C content 42%) suggested that genomic base composition is indeed a driver of the mutational spectrum (Supplement Text 1).

The phenomenon of two different mutations at the same site in different individuals was recently reported also in *D. melanogaster* (Schrider *et al.* 2013) and plausibly explained by a mutation cluster early in the germline with a subsequent error-prone repair. Our results indicate that this may be a relatively frequent process, meriting future attention.

The here presented experimental set-up combines short term mutation accumulation lines with the information of the parental genotypes and is thus an efficient approach to estimate µ in many organisms even without high quality reference genomes. While identifying a substantial number of mutations, the effort in terms of time (5 generations of about 28 days each) and deep sequencing of ten individuals and the reference pool appeared reasonable. Yet, over such a short period, all mutations can still occur in heterozygous fashion, thus revealing the full mutational spectrum (Table S2). The applied experimental design is furthermore promising to determine the influence of demographic, and environmental factors and/or anthropogenic substances on the evolutionary relevant germline mutation rate.

## Acknowledgements

This work was supported by the Deutsche Forschungsgemeinschaft (grant number PF390/8-1). We are grateful to Dennis Lüders for Sanger sequencing and Andreas Wieser for statistical support.

## Author contributions

MP and AMO designed the study, AMO carried out the experiments, MP performed the bioinformatics analyses, MP and AMO drafted the manuscript.

## Data accessibility

Data are deposited at ENA (PRJEB18039).

